# Cohesin guides homology search during DNA repair via loops and sister chromatid linkages

**DOI:** 10.1101/2025.02.10.637359

**Authors:** Federico Teloni, Zsuzsanna Takacs, Michael Mitter, Christoph C. H. Langer, Inès Prlesi, Thomas L. Steinacker, Vincent P. Reuter, Dmitry Mylarshchikov, Daniel W. Gerlich

## Abstract

Accurate repair of DNA double-strand breaks (DSBs) is essential for genome stability, and defective repair underlies diseases such as cancer. Homologous recombination uses an intact homologous sequence to faithfully restore damaged DNA, yet how broken DNA ends find homologous sites in a genome containing billions of non-homologous bases remains unclear. Here, we introduce sister-pore-C, a high-resolution method for mapping intra- and trans-molecular interactions in replicated chromosomes. We show that DSBs reshape chromosome architecture by recruiting two functionally distinct pools of cohesin. Loop-forming cohesin accumulates across a megabase-scale domain to control homology sampling within topologically associating domains (TADs) surrounding the break site, while cohesive cohesin concentrates at the break site to tether broken ends to the sister chromatid. This dual mechanism restricts the homology search space, highlighting how chromosome conformation helps preserve genomic integrity.

---

DNA double-strand breaks (DSBs) are among the most dangerous threats to genome integrity, as incorrect repair can result in chromosomal loss or translocations associated with cancer (*1*). The most accurate mechanism of DSB repair is homologous recombination, which restores the damaged sequence from an intact homologous DNA template (*2, 3*). However, this process requires broken DNA ends to locate a homologous sequence within a genome dominated by non-homologous DNA, creating a substantial kinetic challenge (*4–7*). In eukaryotes, homology search is mediated by nucleoprotein filaments formed by RAD51 protein on single-stranded DNA adjacent to the DSB (*2, 3, 6, 8*). While the interactions between RAD51 filaments and DNA are well-characterized at the molecular scale (*9*), it remains unclear how these filaments navigate the dense, folded chromatin architecture of the nucleus (*10–14*).

DSBs can be repaired by recombination with either the replicated sister chromatid or a homologous chromosome, consistent with a model in which RAD51 filaments scan the entire genome (*4, 6, 7*). Indeed, RAD51 filaments can move throughout the entire cell nucleus in budding yeast (*15*). However, in somatic cells, homologous recombination primarily uses the sister chromatid as a repair template (*2, 6, 7, 16, 17*). This preference is attributed to the formation of cohesin-mediated linkages between sister chromatids, referred to as cohesion (*18–20*), which is essential for homology-directed repair (*21–24*). An alternative model suggests that DSBs recombine with pre-paired molecules rather than scanning the genome (*5*). Yet, in human cells, sister loci are often far apart (*24–27*), and sister chromatids can be locally misaligned by hundreds of kilobases (*27–29*). It remains unclear how broken DNA ends identify homology on sister chromatids despite this substantial misalignment.

Beyond linking sister chromatids, cohesin organizes chromosomes into loops (*30–33*) via motor-driven DNA extrusion (*34, 35*). In vertebrates, cohesin-mediated loop extrusion is restricted by the DNA-binding protein CTCF (*36–38*), thereby structuring the genome into topologically associating domains (TADs) (*39–41*). Cohesin-mediated looping also promotes the dealignment of sister chromatids, especially within TAD bodies (*27, 28*). Yet, the impact of TADs and the dynamic interplay between cohesive- and loop-forming pools of cohesin on homology search remains unknown.

To elucidate how cohesin-regulated chromosome architectures influence DSB repair, we developed methods to map the homology search space in human somatic cells, both within and across sister chromatids. Combining these approaches with targeted degradation of cohesin regulators, we show how loop-forming and cohesive cohesin cooperatively guide homology search.

## Site-specific induction of homology-directed DSB repair

Investigating how chromosome conformation affects homology-directed DNA repair requires precise temporal and positional control of DSBs. To achieve this, we used an inducible Cas9 transgene and transfected a synthetic single-guide RNA (sgRNA) targeting multiple genomic sites (*42*). We chose HeLa cells as a model system because of their proficiency in homology-directed repair and the availability of genome-edited clones enabling targeted degradation of cohesin regulators (*27, 28*). Immunofluorescence staining of the DSB repair factor tumor suppressor p53-binding protein 1 (53BP1) showed robust DSB repair foci about 6 h after sgRNA transfection (Fig. S1A, B).

To induce DSBs specifically during S/G2-phase—when homology-directed repair is active (*43, 44*)—we synchronized cells at the G1/S boundary and released them into S-phase, followed by sgRNA transfection (Fig. S1C-F). By using chromatin immunoprecipitation followed by sequencing (ChIP-seq) with an anti-γH2A.X antibody, we detected DSBs across the genome 6 h after sgRNA transfection, at which point 93% of cells had reached G2 (Fig. S1F). This approach showed robust γH2A.X accumulation at 23 genomic loci (Fig. S1G-I), which we selected for further analysis. We thus established a system for site-specific DSB induction at a cell cycle stage proficient for homology-directed repair.

DSBs can be repaired by either non-homologous end joining or homology-directed repair (*43, 44*). To determine which pathway predominantly repairs Cas9-induced DSBs in G2-synchronized HeLa cells, we visualized nine sgRNA target loci via fluorescence in situ hybridization (FISH), combined with immunofluorescence detection of 53BP1 (marking DSBs) and RAD51 (marking homology-directed repair). Without sgRNA, 53BP1-positive FISH spots ranged from 3% to 6% across target loci, rising to 40–90% with sgRNA transfection (Fig. 1A–F)—confirming the specificity and efficiency of Cas9-mediated DSB induction. Notably, most 53BP1-positive sites also co-localized with RAD51, ranging from 73% to 86% across sgRNA target loci (Fig. 1G). Hence, in G2-synchronized HeLa cells, most Cas9-induced DSBs are repaired by homologous recombination. This experimental system thus provides a robust platform for investigating how chromosome conformation influences homology-directed DNA repair.

**Fig. 1.**
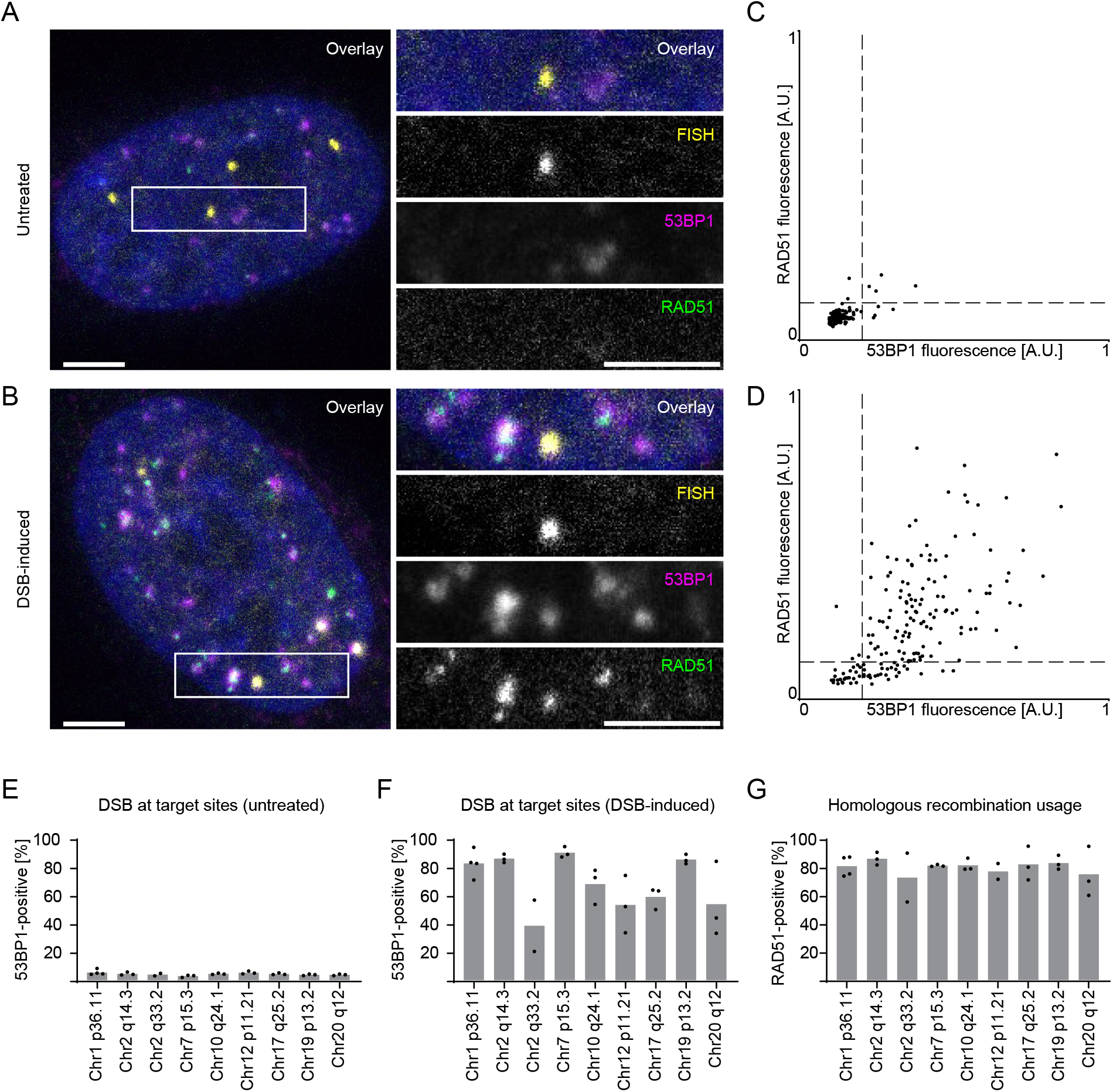
Site-specific induction of homology-directed DSB repair. (**A, B**) Immunofluorescence staining of the general DSB repair factor 53BP1 and the homology-directed-repair-specific factor RAD51 combined with FISH at an sgRNA target locus (chr19 p13.2) in untreated (A) or sgRNA-transfected (DSB-induced) (B) G2-synchronized HeLa cells. Scale bar, 5 µm. (**C, D**) Quantification of 53BP1 and RAD51 fluorescence intensities at the FISH-labeled locus in untreated (C) or DSB-induced (D) cells. Each dot represents one locus; dashed lines indicate 2 S.D. above the mean of the untreated control. (**E, F**) Percentage of FISH-labeled loci positive for 53BP1 in untreated (E) or DSB-induced (F) cells, measured at nine sgRNA target loci. Each dot is the mean from an independent experiment; bars indicate the average of all experiments. (**G**) Fraction of 53BP1-positive loci that also contain RAD51 after sgRNA transfection at the same nine loci. Dots are mean values per experiment; bars indicate overall mean.

## Loop-forming cohesin controls search range

Genomic interactions are normally constrained within local neighborhoods of TADs (*10–14, 31, 45*). However, given the high intrinsic dynamics of RAD51 filaments (*15*), it remains unclear how these architectural constraints affect homology search. To address this, we performed RAD51 ChIP-seq using a library preparation that captures interactions between single-stranded RAD51 nucleoprotein filaments and double-stranded DNA (*46*). Published Hi-C data from G2-synchronized cells (*28*) provided a reference for local chromosome architecture near DSBs (Fig. 2A; S2A). RAD51 ChIP-seq signal accumulated predominantly near sgRNA target sites (Fig. 2B-D; S2A, B), indicating a bias of RAD51 filament sampling toward local genomic neighborhoods rather than uniform sampling of the genome. The RAD51 distribution was often asymmetric relative to sgRNA target sites and was confined by TAD boundaries. Average distributions relative to TAD boundaries near DSBs revealed RAD51 accumulation at TAD boundaries and a sharp decay towards the side facing away from the DSB (Fig. 2E, F). Thus, RAD51 filaments primarily search for homology within TAD-constrained neighborhoods of the break site.

**Fig. 2.**
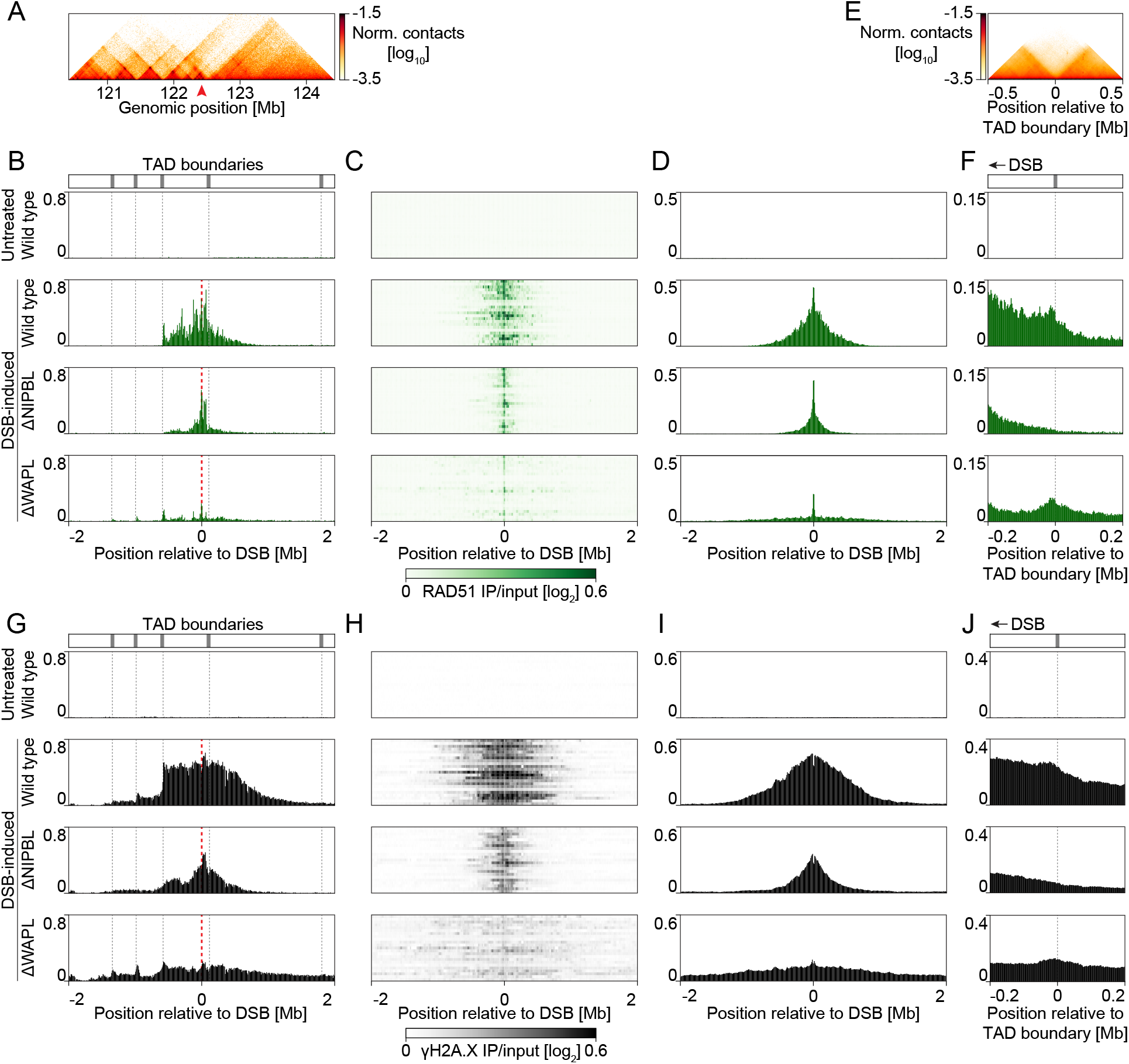
Loop-forming cohesin controls the homology search range. (**A**) Hi-C interaction matrix (*28*) around an sgRNA target locus (chr2 q14.3) in untreated, G2-synchronized cells. The red arrowhead marks the sgRNA target site. (**B**) RAD51 ChIP-seq tracks at the locus shown in (A) for untreated or DSB-induced cells (wild type or with auxin-induced depletion of NIPBL or WAPL). Gray dotted lines indicate TAD boundaries; the red dashed line marks the DSB site. Bin size: 10 kb. (**C, D**) Stacked line profiles (C) and average pileups of RAD51 signals at 23 sgRNA target loci under the conditions in (B). Bin size: 10 kb. (**E**) Average Hi-C interaction profile at TAD boundaries (*28*) surrounding the same 23 loci. (**F**) RAD51 ChIP-seq signal aligned to TAD boundaries from (E), with DSBs positioned at the left. Bin size: 2.5 kb. (**G**–**J**) γH2A.X ChIP-seq tracks (G), stacked line profiles (H), average pileups (I), and TAD-boundary-aligned pileups (J) under conditions in (B). All ChIP-seq signals are shown as log_2_(IP/input) and represent merged data from n = 2 independent experiments.

Because loop-forming cohesin complexes regulate genomic interactions in TADs (*30–33*), we next examined the role of loop-forming cohesin in homology sampling.

Depletion of the cohesin loading and loop-extrusion factor NIPBL (*34, 35, 47*) using an auxin-inducible degron (*27*) (Fig. S2D) substantially reduced the RAD51 ChIP-seq signal spread and abolished RAD51 accumulation at TAD boundaries (Fig. 2B–F). Hence, loop-forming cohesin is critical for defining the genomic range of homology search. Cohesin forms loops on chromatin until it is released by WAPL protein (*48–50*). Depleting WAPL increases loop size and promotes bypass of TAD boundaries (*33, 51*). Because loop size directly influences how DNA segments encounter one another, we investigated the effect of auxin-induced degradation of WAPL (*27*) (Fig. S2E). We observed much less RAD51 signal near DSBs, but a broader spread across multiple TAD boundaries (Fig. 2B-D; S2E). While RAD51 still accumulated at TAD boundaries in WAPL-depleted cells (Fig. 2B–F; S2A, B), it no longer decayed sharply across the boundary (Fig. 2F). Thus, enhanced cohesin-mediated looping expands the homology search range but reduces the likelihood of sampling sequences immediately adjacent to the break site.

DSB repair domains are marked by ATM-mediated γH2A.X phosphorylation, whereby cohesin and TAD boundaries have been proposed to regulate its spread (*52, 53*). To determine how the RAD51 homology search relates to DSB repair domains, we performed γH2A.X ChIP-seq in G2-phase cells following Cas9-mediated DSB induction. Although TAD boundaries constrained γH2A.X spreading, its distribution was broader than the RAD51 signal (Fig. 2G-J; S2C). Consistent with the RAD51 results, NIPBL depletion substantially narrowed γH2A.X spreading, whereas WAPL depletion broadened it (Fig. 2G-J; S2C), corroborating a key role of cohesin-mediated looping in spreading this epigenetic mark. Homology search thus does not extend beyond genomic regions marked by γH2A.X. Collectively, these findings demonstrate that cohesin-mediated loops and TAD boundaries control the genomic space sampled during homology-directed repair.

## Cohesion directs homology sampling across sister chromatids

DSBs recruit cohesin (*53–57*) and locally increase DNA looping (*53*). As enhanced looping promotes separation and local misalignment of undamaged sister chromatids (*27*), this raises the question how broken DNA ends maintain access to the opposing sister chromatid. In budding yeast, DSBs induce cohesion genome-wide (*54, 58, 59*), implying that cohesion remodeling may compensate for chromosome misalignment. However, whether DSBs similarly alter cohesion in human cells remains unclear.

To investigate how DSBs affect chromosome organization, we aimed to map chromatin contacts both within and across sister chromatids. Conventional chromosome conformation capture techniques (*60, 61*) do not distinguish intra-chromatid (cis-sister) from inter-chromatid (trans-sister) contacts. Although sister-chromatid-sensitive Hi-C (scsHi-C) addresses this limitation by labeling sister chromatids with the nucleotide analogue 4-thio-thymidine (*28, 62*), its low labeling efficiency limits resolution. An alternative method uses bromodeoxyuridine (BrdU) for labeling (*63*) but is limited by low sister-chromatid-specificity (*64*). To increase the resolution of sister-chromatid-specific conformation mapping, we developed sister-pore-C (Fig. 3A). This method is based on labeling distinct DNA strands in each sister chromatid by culturing synchronized cells for one S-phase in medium containing BrdU. Combining BrdU detection via Oxford Nanopore sequencing (*65–67*) with Nanopore-based Pore-C (*68*) enables discrimination of cis-sister from trans-sister contacts by assigning DNA-strand orientation and BrdU-label status. This novel pipeline yields multi-fragment concatemers, which increases the number of sister-specific contacts per sequenced read.

**Fig. 3.**
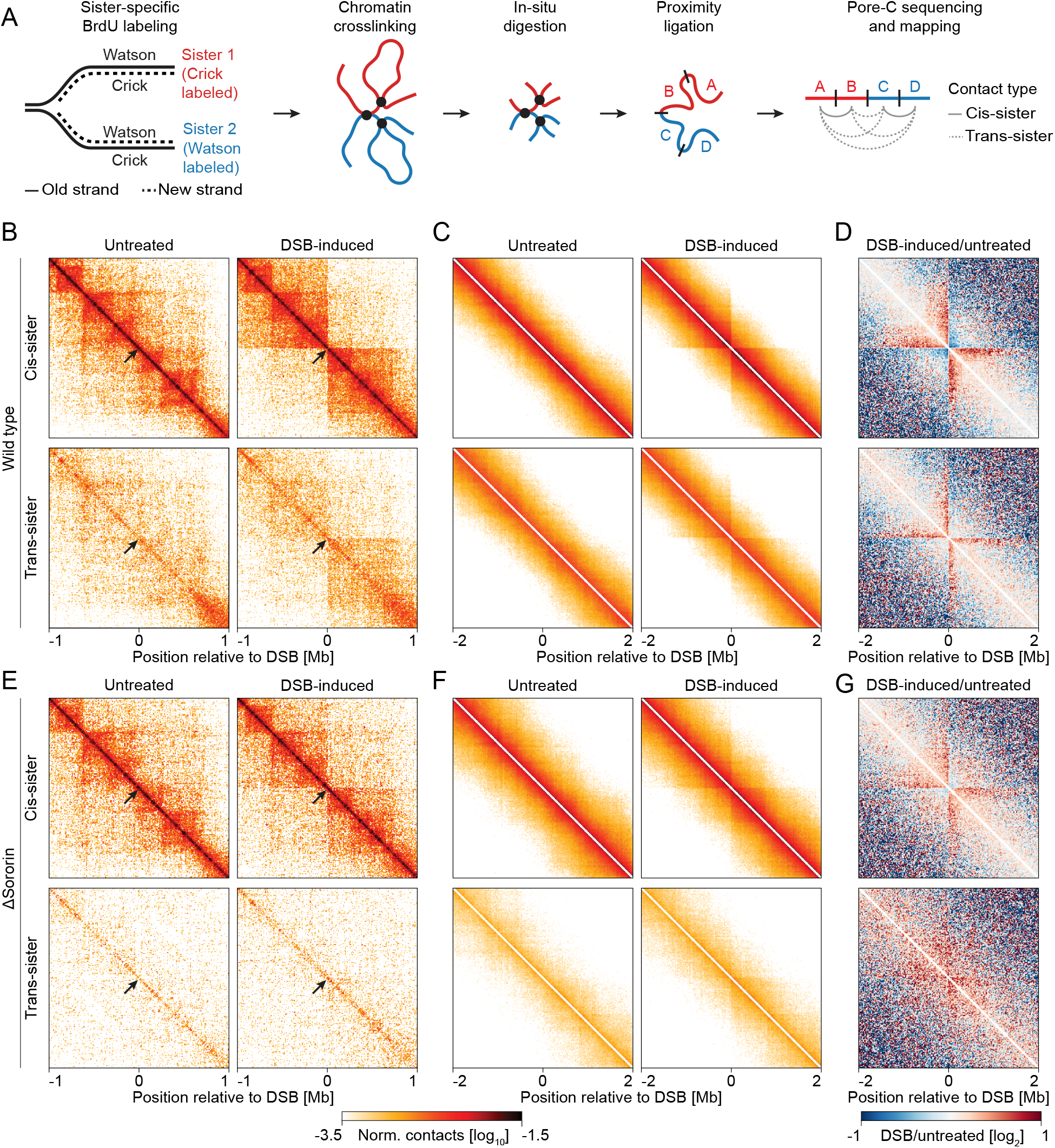
Cohesion directs homology sampling across sister chromatids. (**A**) Schematic of sister-pore-C. Synchronized cells are cultured with BrdU in one S-phase, then crosslinked in G2, followed by in situ digestion and proximity ligation. Nanopore sequencing identifies BrdU labeling and DNA strand orientation, for assignment of each fragment to a sister chromatid and classifying contacts as cis-sister or trans-sister. (**B**) Sister-pore-C interaction matrices of a target locus (chr2 q33.2) in untreated or DSB-induced G2 cells. Black arrows mark the DSB site. Top: cis-sister contacts; bottom: trans-sister. Bin size: 10 kb. (**C, D**) Average pileups (C) and log^2^(DSB-induced/untreated) ratios (D) for cis- and trans-sister contacts across 23 loci (n = 3). Bin size: 20 kb. (**E**–**G**) Sister-pore-C interaction matrices (E) and corresponding pileups (F, G) for Sororin-depleted G2 cells. Bin size: 10 kb (E) and 20 kb (F, G). Data are merged from n = 3 (B–D) and n = 2 (E–G) independent experiments.

We first assessed BrdU incorporation and toxicity in HeLa cells. Culturing cells with 40 µM BrdU for 24 h did not impair proliferation or increase DNA damage (Fig. S3A-C). Mass spectrometry showed BrdU replaced 49% of thymidine in genomic DNA, a 19-fold increase in labeling density compared to scsHi-C (Fig. S3D) (*28, 62*). For sister-pore-C analysis, we released G1/S-synchronized cells into BrdU-containing medium, and cross-linked chromatin in G2 for pore-C analysis (Fig. 3A). Using the DNAscent neural network to detect BrdU (*65, 69*), we classified each DNA segment as labeled or unlabeled; controls from unlabeled cells and from cells grown five days in the presence of BrdU confirmed accurate classification (Fig. S3E). We then categorized each pairwise combination between read segments as cis-sister or trans-sister contact based on the BrdU labeling status and strand orientation. This approach yielded three sister-specific contacts per mapped read segment in average, a 50-fold increase over the previous scsHi-C method (*28, 62*). Sister-pore-C thus allows high-resolution analysis of chromosomal interactions during homology-directed DNA repair.

To validate sister-pore-C, we compared sister-pore-C contact maps in G2-synchronized cells with previously published scsHi-C data (*28*) and evaluated auxin-induced Sororin depletion, which disrupts cohesion without affecting cohesin loops (*28, 70, 71*). Sister-pore-C maps closely matched scsHi-C results but required 50-fold fewer read segments to achieve comparable contact density and resolution (Fig. S4A-C). Sororin depletion selectively reduced trans-sister contacts without affecting cis-sister contacts (Fig. S4D), confirming accurate sister assignment. Thus, sister-pore-C reliably captures cis- and trans-sister contacts, confirming its suitability for studying cohesin-regulated chromosome interactions.

To determine the effects of DSBs on chromosome architecture, we compared sister-pore-C maps of DSB-containing and undamaged chromosomes. We induced DSBs in S/G2-synchronized cells using Cas9 (Fig. S5A) and generated cis- and trans-sister contact maps to examine local and genome-wide changes. DSBs increased cis-sister contacts in their vicinity (Fig. 3B-D; S5B), consistent with conventional Hi-C analysis of DSBs in unsynchronized cells (*53*). Trans-sister maps revealed a prominent contact stripe extending over about 1 Mb from the DSB (Fig. 3B-D; S5B), indicating that DSBs enhance interactions between broken DNA ends and the opposing sister chromatid. Genome-wide average contact probabilities, however, were unaffected by DSBs (Fig. S5C, D). Thus, DSBs locally remodel chromosome architecture during homology-directed repair, promoting increased looping in their vicinity and targeted interactions between broken DNA ends and the sister chromatid.

To investigate whether the DSB-induced trans-sister interactions are mediated by cohesion, we applied sister-pore-C to Sororin-depleted cells and compared cis- and trans-sister contacts around DSBs. Sororin depletion nearly abolished DSB-induced trans-sister contact stripes, whereas cis-sister contacts remained elevated around DSBs (Fig. 3E-G; S5B, E). Thus, rather than increasing sister chromatid pairing genome wide, as in budding yeast (*58, 59*), DSBs specifically promote cohesion-dependent interactions between broken DNA ends and the opposing sister chromatid in human cells.

## Two cohesin pools guide homology search

The remodeling of chromosome architecture by DSBs raises the question of how cohesin-regulated loops and trans-sister contacts are spatially organized relative to one another. Profiling DSB-induced contact changes along the genome showed increased looping over approximately one Mb relative to sgRNA target sites, whereas trans-sister contacts concentrated in a much narrower region (Fig. 4A-D).

**Fig. 4.**
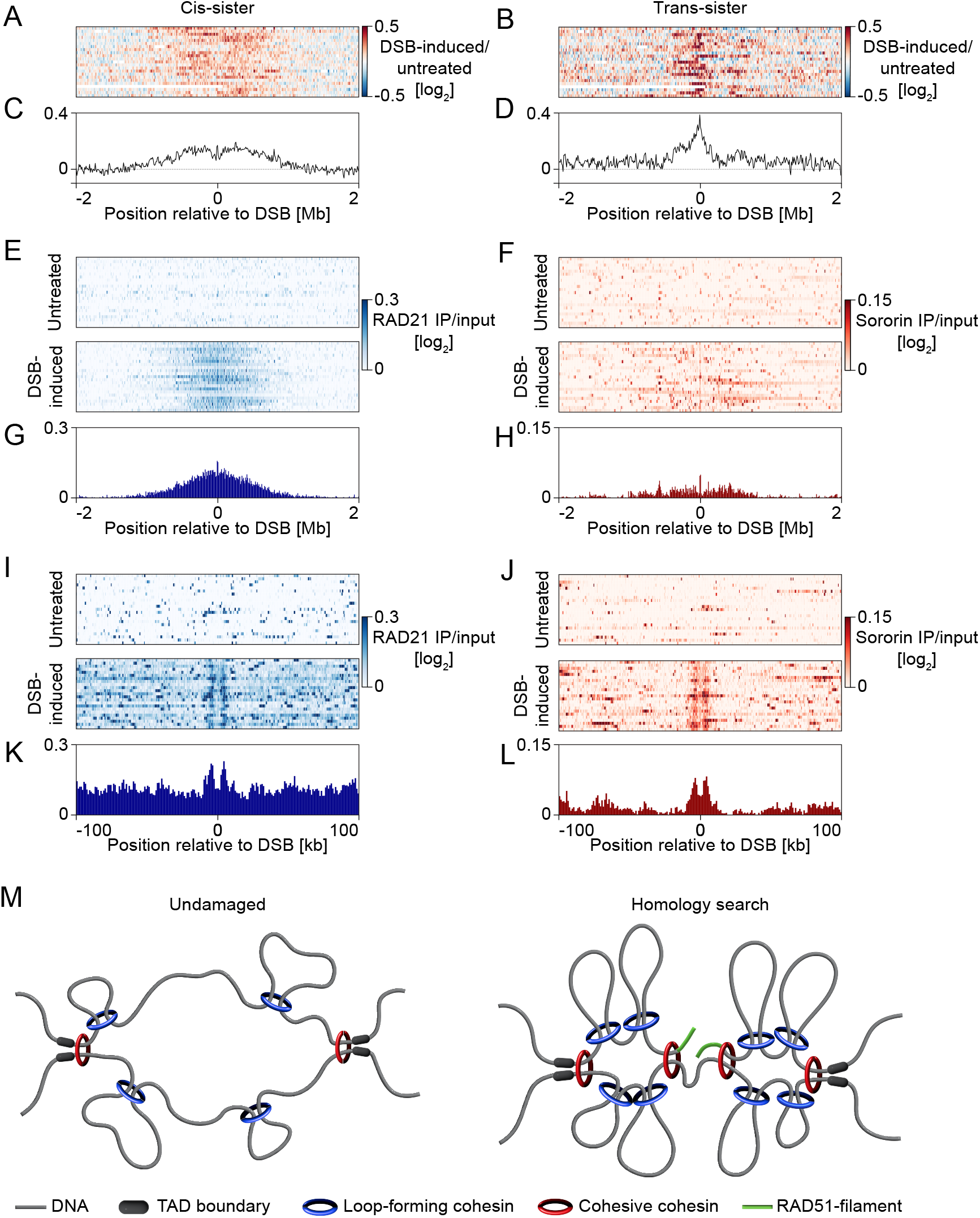
Distinct roles for loop-forming and cohesive cohesin in homology search. (**A, B**) Stacked line plots of log^2^(DSB-induced/ untreated) for cis-sister (A) and trans-sister (B) contacts at 23 loci, evaluated over 30–1000 kb. Bin size: 10 kb. Data are merged from n = 3 independent experiments. (**C, D**) Average pileups of the data in (A) and (B). Bin size: 10 kb. (**E, F**) Stacked line profiles of RAD21 and Sororin (F) ChIP-seq signals at the same 23 loci in untreated or DSB-induced G2 cells. Bin size: 10 kb. (**G, H**) Average ChIP-seq pileups of RAD21 (G) and Sororin (H) in DSB-induced G2 cells, shown as log^2^(IP/input). Bin size: 10 kb. (**I, J**) Higher-resolution stacked profiles of RAD21 (I) and Sororin (J). Bin size: 1 kb. (**K, L**) Corresponding high-resolution pileups of RAD21 (K) and Sororin (L) in DSB-induced G2 cells, shown as log^2^(IP/input). Bin size: 1 kb. All ChIP-seq data are merged from n = 2 independent experiments. Model: DSBs recruit a cohesive clamp that tethers RAD51 filaments near the sister chromatid, while loop-forming cohesin promotes lateral mobility within the TAD. Together, these mechanisms narrow the homology search space, enabling efficient DNA repair.

To further examine how DSBs influence cohesin-regulated chromosome architecture, we mapped the distribution of the core cohesin subunit RAD21, which marks both loop-forming and cohesive cohesin, and the cohesion-specific cofactor Sororin using ChIP-seq. RAD21 showed a substantial accumulation across a ~1 Mb region surrounding the DSB, consistent with the observed increase in looping (Fig. 4A, C, E, G). Sororin, by contrast, showed very little enrichment across this broad domain (Fig. 4F, H) but was sharply concentrated in the immediate vicinity of DSBs, overlapping with RAD21 at the break site (Fig. 4I-L). These results suggest that DSBs recruit loop-forming and cohesive cohesin to distinct architectural roles. While loop-forming cohesin broadly spreads across the 1 Mb domain, cohesive cohesin concentrates near the DSB to mediate trans-sister interactions.

## Discussion

Here we reveal how cohesin-regulated chromosome architecture governs homology search in somatic cells. Our findings support a model where DSBs trigger the formation of a cohesive clamp at broken DNA ends to facilitate inter-sister homology search, while increased looping by cohesin controls lateral sampling within the local TAD (Fig. 4M). This dual mechanism compensates for local sister-chromatid misalignment prior to DNA damage and substantially narrows the homology search space compared to a genome-wide scan, providing a major kinetic advantage.

Positioning cohesive cohesin near RAD51 filaments may involve lateral sliding or de novo loading, potentially mediated by Smc5/6 (*56*) or the MRN complex (*55, 72*). Meanwhile, loop-forming cohesin may be loaded to more distal sites by γH2A.X (*55*). The controlled DSB induction assay described here paves the way for future dissection of the molecular events coordinating these distinct cohesin pools during homology-directed repair.

Beyond its role in maintaining genomic integrity, DNA recombination is a fundamental driver of genetic diversity, promoting antibody diversification and the exchange of genetic material between maternal and paternal genomes during meiosis. These events involve distinct topologies: antibody diversification is mediated by cohesin-regulated intramolecular loops (*73, 74*), whereas meiosis primarily depends on recombination between paired homologous chromosomes (*75*). By enabling simultaneous, genome-wide profiling of both intra- and intermolecular interactions, Sister-Pore-C will provide unprecedented insights into the role of chromosome topology in governing genome evolution and stability across diverse biological processes.

## Acknowledgements

The authors thank the IMBA/IMP/GMI BioOptics and Molecular Biology Service and the Vienna BioCenter Next Generation Sequencing and Metabolomics facilities for technical support, J. Ellenberg and K.S. Beckwith for technical advice on RASER-FISH, and J.-M. Peters for providing Sororin-AID and WAPL-AID HeLa cell lines, R. Medema for providing the iCut plasmid, E. Ogris for the production of the α-Sororin antibody, and I. Patten, M.M. Alaabo, and T. Hidaka for comments on the manuscript. **Funding:** Research in the laboratory of DWG has been supported by the Austrian Academy of Sciences, the Vienna Science and Technology Fund (WWTF; project LS19-001), and the European Research Council (ERC) under the European Union’s Horizon 2020 research and innovation programme (grant agreement no. 101019039). DWG is also an adjunct professor at the Medical University of Vienna. FT has received fellowships from the Swiss National Science Foundation (P2ZHP3_191408) and from the European Union’s Horizon 2020 research and innovation programme (Marie Skłodowska-Curie Individual Fellowship 101022896 – DSB Architect). TS has received fellowships from the European Union’s Framework Programme for Research and Innovation Horizon 2020 (Marie Curie Skłodowska Grant Agreement Nr. 847548), the European Molecular Biology Organization (ALTF 866-2022), and from the European Union’s Horizon Europe research and innovation programme (Marie Skłodowska-Curie Individual Fellowship 101103258 – MicroChrom). ZT has received a Hertha Firnberg Programme fellowship of the FWF (T 1246). MM has received a PhD fellowship from the Boehringer Ingelheim Fonds. **Author contributions:** Conceptualization: FT, DWG. Methodology: FT (DSB induction assay); ZT, MM, CCHL (sister-pore-C); TS, VPR (FISH). Investigation: IP (Fig. 1), DM (Fig. 4A, B, S5C, D), FT (all other data). Visualization: IP (Fig. 1), CCHL (Fig. S4), DM (Fig. 4A, B, S5C, D), FT (all other Figs.). Funding acquisition: DWG, FT, ZT, MM, TS. Supervision: DWG. Writing – original draft: DWG, FT. Writing – review & editing: All authors. **Competing interests:** The authors declare no competing interests. **Data and material availability:** All data and code will be made available in public repositories upon publication of the manuscript. All materials will be made available upon request to the corresponding authors.

## Materials and methods

### Cell culture

All cell lines were cultured under sterile conditions in a humidified incubator at 37 °C with 5% CO2. Regular testing confirmed they were free of mycoplasma contamination. The parental HeLa cell line (Kyoto strain) was obtained from S. Narumiya (Kyoto University, Japan) and validated using the Multiplex Human Cell Line Authentication (MCA) test. All lines were grown in Dulbecco’s Modified Eagle’s Medium (DMEM; IMP/ IMBA/GMI Molecular Biology Service/Media Kitchen) supplemented with 10% (v/v) fetal bovine serum (FBS; Gibco, 10270-106), 1% (v/v) GlutaMAX (Invitrogen, 35050038), and 1% (v/v) penicillin-streptomycin (Sigma-Aldrich, P0781), hereafter referred to as wild-type medium. For Cas9-dox cell lines, the medium additionally contained 0.5 mg/ml G418 (Invitrogen, 11811-031). Sororin-AID and WAPL-AID lines were maintained with 0.5 μg/ml puromycin (Calbiochem, 540411), and NIPBL-AID lines were grown with 6 μg/ml blasticidin S (Thermo Fisher Scientific, A1113903). All cell lines used in this study are listed in Supplementary Table 1.

### Generation of Cas9-dox cell lines

300,000 HeLa cells (wild-type parental, Sororin-AID (28), WAPL-AID, or NIPBL-AID (27)) were seeded into individual wells of a 6-well plate in wild-type medium. The next day, cells were transfected with a total of 0.66 μg plasmid DNA encoding a humanized Streptococcus pyogenes Cas9 (gift from R. Medema) fused to FKBP12^F36V^-5xGly-3xFLAG at the N-terminus, under a tetracycline (Tet)-inducible promoter in a transposon array, along with 0.33 μg Super PiggyBac Transposase plasmid DNA (Systembio, PB210PA-1). Transfections were performed in 100 μl OptiMEM (Thermo Fisher Scientific, 31985070) using 4 μl polyethyleneimine (Polysciences Europe, 24765-2). 24 hours later, cells were harvested by trypsinization (Thermo Fisher Scientific, 25200056) and transferred to 15-cm dishes in wild-type medium. After another 24 hours, 0.5 mg/ml G418 and the respective selection markers for each cell line (see Supplementary Table 1) were added. Cells were cultured under selection for two weeks to generate polyclonal populations expressing FKBP12^F36V^-5xGly-3xFLAG-Cas9 upon induction with 0.2 μg/ml doxycycline (Sigma-Aldrich, D9891). For Cas9-dox WT and NIPBL-AID lines, single-cell dilution was performed in 96-well plates. Clonal populations were expanded for 11 days under the relevant selection antibiotics. All lines were tested to confirm Cas9 expression following doxycycline induction.

### sgRNA transfection

Thes gRNA sequence (5′-CAGACAGGCCCAGATTGAGG-3′) used in this study was obtained from (42). Transfections were performed with TrueGuide sgRNA (Thermo Fisher Scientific) and Lipofectamine RNAiMAX (Thermo Fisher Scientific, 13778150) for up to 8 hours. Unless otherwise noted, the final sgRNA concentration was 27.5 nM.

### Immunofluorescence

For immunofluorescence (IF), cells were washed twice with phosphate-buffered saline (PBS) and fixed with 4% formaldehyde (Thermo Fisher Scientific, 28906) in PBS for 5 minutes at room temperature (RT), if not stated otherwise. Samples were then permeabilized with 0.2% Triton X-100 (Sigma-Aldrich, 327371000) in PBS for 5 minutes. After blocking with filtered 2% bovine serum albumin (BSA; Sigma-Aldrich, A7030) in PBS for 15 minutes at RT, cells were incubated with primary antibodies diluted in 2% BSA/PBS for 2 hours at RT. Following three washes in PBS, cells were incubated with fluorescently labeled secondary antibodies for 1 hour at RT in the dark. Nuclei were stained with 0.5 μg/ml 4′,6-diamidino-2-phenylindole (DAPI; Thermo Fisher Scientific, 62248) for 10 minutes, washed briefly in PBS, and mounted with ProLong Diamond Antifade Mountant (Thermo Fisher Scientific, P36970). Primary antibodies used were α-53BP1 (1:1000; Millipore, MAB3802), α-γH2A.X (phospho S139) (1:1000; Abcam, ab81299), and α-RAD51 (1:1000; Abcam, ab176458). Secondary antibodies were α-mouse IgG Alexa Fluor 568 (1:1000; Molecular Probes, A11004) and α-rabbit IgG Alexa Fluor 488 (1:1000; Molecular Probes, A21206).

### sgRNA transfection time-course

Cas9-dox WT HeLa Kyoto cells were seeded onto 12-mm glass coverslips at ~30% confluence and grown for 24 hours in wild-type DMEM supplemented with 0.2 μg/ml doxycycline (Sigma-Aldrich, D9891) to induce Cas9 expression. Cells were transfected with sgRNA to induce DSBs at target loci and harvested every 2 hours up to 8 hours post-transfection. Samples were processed to detect 53BP1 (see Immunofluorescence). Images were acquired on an Olympus ScanR Screening System (Olympus IX83 microscope, 40× objective, 0.75 NA) and analyzed with Olympus ScanR Image Analysis Software (v3.2.0) to quantify 53BP1 foci formation using a spot detection module (76).

### Cell synchronization to G2-phase and DSB induction

Cells were seeded in wild-type medium and allowed to attach for 3 hours before adding 2 mM thymidine (Sigma-Aldrich, T1895) to synchronize them at the G1/S boundary. After 16 hours, cells were washed twice with pre-warmed PBS and released into wild-type medium for 8 hours. Cells were then treated with 3 μg/ml aphidicolin (Sigma-Aldrich, A0781) to synchronize again at the G1/S boundary. Next, cells were washed twice with pre-warmed PBS, released into wild-type medium and collected 9 hours later. For DSB repair experiments, 0.2 μg/ml doxycycline (Sigma-Aldrich, D9891) was added concurrently with aphidicolin to induce Cas9 expression. sgRNA transfection was performed 3 hours after aphidicolin release. For WAPL-AID and NIPBL-AID experiments, 500 μM indole-3-acetic acid (auxin; Sigma-Aldrich, I5148) and 1 μM 5-Ph-indole-3-acetic acid (5-Ph-auxin; Bio Academia, 30-003) were added concurrently with the release into wild-type medium.

### Flow cytometry—DNA content analysis and cell cycle stage verification

Flow cytometry was used to verify cell cycle stage. A fraction (5–10%) of harvested cells was washed with PBS and fixed in 70% ethanol (Sigma-Aldrich, 32221) at 4 °C for at least 30 minutes. Fixed cells were permeabilized with 0.25% Triton X-100 (Sigma-Aldrich, 327371000) in PBS for 15 minutes on ice, then incubated with 0.25 μg α-phospho-H3Ser10 (Millipore, 05-806) in 1% BSA/PBS for 1 hour at RT. After washing in 1% BSA/PBS, samples were incubated for 30 minutes at RT in the dark with α-mouse IgG Alexa Fluor 488 (1:300; Molecular Probes, A11001). DNA content was quantified by staining with 200 μg/ml RNase A (Qiagen, 19101) and 50 μg/ml propidium iodide (Sigma-Aldrich, 81845) for 30 minutes in the dark. Data were acquired on a Penteon flow cytometer (Novacyte) and analyzed in FlowJo (v10). FSC-A/SSC-A was used for cell gating, and singlets were identified using FSC-A/FSC-H and PI-A/PI-H. H3S10P (FITC channel) versus DNA content (PI channel) provided cell cycle profiles.

### Chromatin immunoprecipitation (ChIP)

Approximately 10 million G2-synchronized HeLa cells (see Cell synchronization to G2-phase and DSB induction) were harvested by washing once with PBS, detaching with trypsin, and collecting in 15-ml Falcon tubes. All centrifugation steps were at 1,100 g for 1 minute unless otherwise specified. Cells were washed once with PBS, fixed in 10 ml of 1% formaldehyde (Thermo Fisher Scientific, 28906) for 10 minutes at RT with gentle rotation, and quenched with 125 mM glycine (Sigma-Aldrich, 50046) for 5 minutes at RT. Cells were pelleted, the supernatant was removed, and pellets were stored at −80 °C. One day prior to processing the samples for ChIP, 320 μl of a 1:1 Affi-Prep Protein A bead suspension (Bio-Rad, 1560005) per sample was pelleted, the supernatant removed, and incubated in 10 volumes of coating buffer (2 mM EDTA, 20 mM Tris-HCl [pH 8.0], 1% Triton X-100, 150 mM NaCl, 1 mM PMSF, 0.1 mg/ml BSA) overnight at 4 °C with gentle rotation. Beads were washed three times in chilled dilution buffer (2 mM EDTA, 20 mM Tris-HCl [pH 8.0], 1% Triton X-100, 150 mM NaCl, 1 mM PMSF) and stored as a 1:1 suspension in dilution buffer at 4 °C. Unless otherwise noted, all sample processing steps were performed on ice. Cell pellets were resuspended in 1 ml freshly prepared, chilled lysis buffer (10 mM EDTA, 50 mM Tris-HCl [pH 8.0], 1% SDS, 1 mM PMSF, cOmplete mini protease inhibitor cocktail), pelleted at 1,700 g for 2 minutes at 4 °C, and resuspended in 700 μl lysis buffer. Chromatin was sonicated in a Bioruptor Plus (Diagenode) for 12 cycles (30 s ON, 30 s OFF) at high intensity at 4 °C. After sonication, 10 μl was saved as input; the remaining chromatin was diluted 10-fold (7 ml) in dilution buffer. Pre-clearing was performed by incubating samples with 160 μl 1:1 blocked Protein A beads for 1 hour at 4 °C, followed by centrifugation. Sonicated chromatin was quantified using a DS-11 spectrophotomer (DeNovix) and incubated overnight at 4 °C with one of the following antibodies: ≥25 μg chromatin with 4 μg α-RAD51 (Abcam, ab176458), ≥25 μg chromatin with 2 μg α-γH2A.X (Abcam, ab81299), ≥50 μg chromatin with 10 μg α-RAD21 (Abcam, ab992), ≥100 μg chromatin with 10 μg α-Sororin (Abcam, ab192237). Immunoprecipitation was performed the next day by adding 160 μl 1:1 blocked Protein A beads for 3 hours at 4 °C with gentle rotation. Beads were washed twice with 800 μl of each of the following pre-chilled buffers: washing buffer 1 (2 mM EDTA, 20 mM Tris-HCl [pH 8.0], 1% Triton X-100, 150 mM NaCl, 0.1% SDS, 1 mM PMSF), washing buffer 2 (2 mM EDTA, 20 mM Tris-HCl [pH 8.0], 1% Triton X-100, 500 mM NaCl, 0.1% SDS, 1 mM PMSF), washing buffer 3 (2 mM EDTA, 10 mM Tris-HCl [pH 8.0], 250 mM LiCl, 0.5% NP-40, 0.5% sodium deoxycholate), and TE buffer (1 mM EDTA, 10 mM Tris-HCl [pH 8.0]). After the final TE wash, beads were resuspended in 200 μl pre-warmed elution buffer (5 mM EDTA, 25 mM Tris-HCl [pH 8.0], 0.5% SDS) and incubated at 65 °C for 20 minutes with shaking (1,200 rpm). The supernatant was collected, and a second elution was performed. Both ChIP and input chromatin were treated with 2 mg/ml RNase A (Qiagen, 19101) and 2 mg/ml Proteinase K (Qiagen, 19133) for 1 hour at 37 °C, followed by overnight de-crosslinking at 65 °C. DNA was purified with the Monarch PCR & DNA Cleanup Kit (NEB, T1030), eluted in 30 μl H2O, and quantified using a Qubit Fluorometer (Thermo Fisher Scientific).

### ChIP-sequencing, read processing, and data analysis

ChIP and input libraries were prepared using the NEBNext Ultra II kit (NEB, E7645S) following the manufacturer’s protocol. Samples with dual unique index barcodes were pooled at equimolar ratios and sequenced on Illumina NextSeq 550 or NovaSeq X platforms, employing 75-bp single-end or 150-bp paired-end read mode, respectively. Library demultiplexing followed Illumina guidelines. Data were analyzed using the nf-core/chipseq pipeline (v1.2.1; https://nf-co.re/chipseq/1.2.1), with parameters “--single_end” and “--hard_trim5 75” to maintain consistent read lengths across different sequencing runs. Reads were aligned to the human genome (hg19) using default settings. After pipeline completion, BAM files for biological replicates of the same sample were merged and re-indexed with *SAMtools* (*77*). Merged files were normalized to counts per million (CPM), and log^2^(ChIP/Input) BigWig files were created using *deepTools* (*78*). These normalized files were used for all ChIP-seq visualizations.

### Karyoplots

R scripts for karyoplots are in the main GitHub repository. We used *karyoploteR* (*79*) and *rtracklayer* (*80*) to visualize sgRNA target sites (Fig. S1G), log^2^(γH2A.X IP/Input) ratios (Fig. S1H), and the selection of target sites (Fig. S1H) over a karyotype plot.

### Selection of sgRNA target locations for genomics analysis

Genomic sites targeted by the sgRNA were identified through γH2A.X ChIP enrichment following DSB induction (see “sgRNA transfection” and “Chromatin immunoprecipitation”). For each predicted sgRNA target (0–2 bp mismatch), the average log^2^(γH2A.X IP/Input) was calculated in a ±25 kb region around the target. Targets with an average log^2^ ≥ 0.2 were designated “enriched” and selected for analysis and are listed in Supplementary Table 2. The DSB on chr22 was excluded because it did not match the sgRNA sequence in hg19.

### Stack-up profiles and average pileups

ChIP-seq protein distributions were visualized through stacked and average pileup profiles generated from BigWig files using the Python *bbi* package (https://github.com/nvictus/python-bbi). NaN values were set to zero to avoid artifacts. The *stackup* function aligned signals across genomic regions, producing heatmaps (stacked profiles) and average signals (pileups). Initial exploration of datasets was performed using HiCognition (*81*), while final plots were created through Jupyter notebooks using *matplotlib* (*82*) and *seaborn* (*83*).

### FISH methodology

FISH probe generation and hybridization followed established protocols (*84, 85*) based on RASER-FISH (*86*) and oligo pool amplification (*87*).

### FISH probe design and amplification

A HeLa Kyoto reference genome (hg19 with SNP correction (*28*)) was used for primary FISH probe design. We selected 9 of the 23 gRNA target sites and designed probes for a 70 kb region surrounding the DSB. To prevent overlap with resected RAD51-coated filaments, the ±5 kb regions flanking the cut site were excluded from the design. OligoMiner (Beliveau et al. 2018) was used to identify 36–42 bp unique target sequences (42 °C annealing temperature [at 50% formamide concentration], 30–70% GC content, 0.25–0.99 LDA stringency, 5–20 k-mer filtering, and a 0.1 secondary structure filter). At least 300 probes covered each DSB site. Each unlabeled primary oligo carried short barcode sequences for binding to fluorescently labeled secondary oligos, as well as flanking primers for oligo pool amplification containing the T7 in vitro transcription (IVT) promoter. Oligo pools were ordered from GenScript (GenTitan™ Oligo Pools). FISH probes were amplified by qPCR using 2× Phusion High-Fidelity PCR Master Mix (Thermo Fisher Scientific, F531), 15 ng oligo pool, 0.5 μM forward and reverse primers, and 1× EvaGreen (Biotium, 31000). Samples were incubated at 98 °C for 3 min, followed by 98 °C for 10 s, 66 °C for 10 s, and 72 °C for 15 s in repeated cycles until near plateau. Amplified DNA was purified using the DNA Clean & Concentrator-25 kit (Zymo Research, D4033) and eluted in 50 μl ultrapure water. For IVT, 1.5 μg PCR product was incubated overnight at 37 °C using the HiScribe T7 Quick High Yield RNA Synthesis Kit (NEB, E2050) with 80 μl NTP buffer mix, 6.25 μl T7 RNA polymerase mix, 6.25 μl RNAsin Plus RNAse inhibitor (Promega, N2615), and ultrapure water to reach 160 μl. 150 μl of unpurified ssRNA was used directly for reverse transcription (RT) at 50 °C for 1 hour with 21 μl of 25 mM dNTP mix (final concentration 1.75 mM), 57 μl of 100 μM forward primer (final concentration 19 μM), 6 μl RNAsin Plus RNAse inhibitor, 6 μl Maxima H Minus Reverse Transcriptase (1,200 U, Thermo Fisher Scientific, EP0753), and 60 μl of 5× RT buffer. RNA was then degraded by 150 μl 0.5 M EDTA and 150 μl 1 M NaOH at 95 °C for 15 minutes. The ssDNA oligo pool was purified with a DNA Clean & Concentrator-100 kit (Zymo Research, D4030), eluted in 200 μl ultrapure water, and its concentration determined by Nanodrop. Fluorescently labeled secondary oligos were synthesized via click chemistry (ClickTech Oligo Link Kit, baseclick) with 3′-C3-azide oligos (Metabion) and an alkyne-modified ATTO647N fluorophore (ATTO-TEC, AD 647N). After purification with two n-butanol washes (Sigma-Aldrich, B7906) or HPLC, labeled oligos were diluted in 1× TE buffer.

### RASER-FISH

HeLa cells were seeded in Ibidi μ-Slide VI 0.5 Glass Bottom slides at 1.5×10^5^ cells/ml (120 µL per channel). Cell synchronization and DSB induction followed the protocol described in “Cell synchronization to G2-phase and DSB induction,” with 40 μM BrdU/BrdC (3:1 ratio; Sigma-Aldrich, B5002; Santa Cruz Biotechnology, sc-284555) added after aphidicolin washout. Cells were fixed at RT for 15 min with 4% PFA (Electron Microscopy Sciences, 15710) in PBS and then quenched for 10 min with 100 mM NH^_4^Cl (Sigma-Aldrich, 326372). Immunofluorescence against 53BP1 and RAD51 was performed as described (see Immuno fluorescence), except fixation and permeabilization were each 15 minutes and the DAPI staining was skipped. A secondary fixation was performed with 1 mM Bis(NHS)PEG5 (Sigma-Aldrich, 803537) in PBS for 30 min at RT, followed by an additional quenching step. Cells were permeabilized for 20 min in PBS with 0.2% Triton-X-100 (Sigma-Aldrich, 327371000) at RT, stained with 0.5 μg/ml DAPI in PBS for 15 min at RT, then UV-treated in a Vilber Bio-Link crosslinker (254 nm, 3.6 J/cm2) to nick BrdU/C-incorporated strands. Nicked strands were degraded with Exonuclease III (1 U/μl; NEB, M0206) at 37 °C for 15 min. Cells were pre-incubated in FISH hybridization buffer (50% (v/v) formamide; Thermo Fisher Scientific, AM9342; 10% (w/v) dextran sulfate; Merck, S4030; 2×SSC; Thermo Fisher Scientific, AM9763) at 37 °C for 1 hour, followed by overnight incubation at 42°C with primary FISH probes (~2 nM/probe) in the same buffer. After two 35 °C washes (5 min each) in 50% (v/v) formamide/2×SSC, cells were washed in 2×SSC+0.2% (v/v) Tween-20 (Sigma-Aldrich, P9416). RNA-DNA hybrids were removed by RNase H (0.05 U/μl; NEB, M0297) at 37 °C for 20 min, followed by 2–3 room-temperature washes in 2×SSC+0.2% Tween-20. Secondary FISH hybridization was performed at RT for 3 min in a buffer containing 5% (w/v) ethylene carbonate (Sigma-Aldrich, E26258) and fluorescent secondary oligos (20 nM) in 2×SSC+0.2% Tween-20. Excess secondary oligos were removed by a 1-min wash at RT with 10% (v/v) formamide in 2×SSC+0.2% Tween-20. Cells were then incubated in 2×SSC+0.2 µg/ml DAPI before imaging. All IF+FISH experiments were imaged on a Zeiss LSM980 confocal system using a 63× NA 1.4 oil DIC Plan-Apochromat objective (ZEN Blue 2020 software v3.7). Images were collected in confocal mode with optimal sectioning (390 nm between z-slices).

### RASER-FISH—Image analysis

Images were processed with an in-house automated pipeline (v0.9.0; https://github.com/gerlichlab/reparafil), except for a small subset of images which were analyzed manually. Prior to pipeline processing, nuclei were segmented based on DAPI staining in Fiji. These nuclear regions (stored as ImageJ .roi files) were used to exclude non-nuclear FISH signals. Fluorescent puncta (FISH spots) were identified using a pixel-intensity threshold approach via the *spotfishing* package (https://github.com/gerlichlab/spotfishing). Immunofluorescence intensities were collected from 10-pixel boxes around each FISH spot centroid in the relevant z-slice. To classify a FISH spot as 53BP1-positive, a signal threshold was set to the mean plus two standard deviations of the 53BP1 intensity in non-targeted controls; this threshold was determined independently for each target site. Spots exceeding the threshold were scored as 53BP1-positive. For these 53BP1-positive spots, homologous recombination was assessed by RAD51 intensity, using the same thresholding approach.

### TAD calling

TAD boundaries from (*28*) were used, obtained by OnTAD (*88*) on a G2 WT Hi-C matrix merged over all replicates (GSM4613674). For the analysis focused on TAD boundaries in Fig. 2E, all boundaries located between 200 and 1000 kb upstream and downstream of the 23 selected genomic loci were included.

### Western Blot

G2-phase synchronized Cas9-dox wild-type, Sororin-AID, WAPL-AID, or NIPBL-AID cell suspensions (0.5 million cells/mL) with or without auxin treatment were prepared for protein analysis by mixing cells with 6× SDS loading buffer containing 10 mM DTT (Roche) and boiling at 95 °C for 5 minutes. Proteins were resolved on NuPAGE Bis-Tris 4–12% gels (for Sororin-AID and WAPL-AID) or NuPAGE Tris-Acetate 3–8% gels (for NIPBL-AID) (Thermo Fisher Scientific, NP0335BOX and EA0378BOX) and transferred to Hybond P 0.45 PVDF membranes (GE Life Sciences). A wild-type control sample was run on each gel to assess target protein degradation. Membranes were probed with α-Sororin (1:50, in-house), α-NIPBL (1:800; Absea, 010702F01), or α-WAPL (1:1000; Santa Cruz, sc-365189).

For loading controls, α-Vinculin (1:10,000; Abcam, ab129002) was used for Sororin-AID and NIPBL-AID samples, and α-Tubulin (1:5,000; Abcam, ab52866) was used for WAPL-AID samples. HRP-conjugated secondary antibodies were goat α-rat (1:2,000; Amersham, NA935), goat α-rabbit (1:5,000; Bio-Rad, 170-6515), and goat α-mouse (1:5,000; Bio-Rad, 1706516). Signal was detected with Bio-Rad Clarity Max ECL (Bio-Rad, 1705062) on a Bio-Rad ChemiDoc imager.

### Cell proliferation and DNA damage assay

HeLa Kyoto cells were seeded onto 12-mm glass coverslips and grown for 24 hours. Cells were then treated with various concentrations of 5-bromo-2′-deoxyuridine (BrdU; 10–100 μM; Life Technologies, B23151) and 50 μM etoposide (Sigma-Aldrich, E1383) for an additional 24 hours. Samples were processed to detect γH2A.X (see Immunofluorescence). Images were acquired on an Olympus ScanR Screening System (Olympus IX83 microscope, 40× objective, 0.75 NA) and analyzed in Olympus ScanR Image Analysis Software (v3.2.0) to quantify nuclei and immunofluorescence intensity (*76*). Fold increase in cell number was calculated by comparing the total segmented nuclei count to the initial seeding density.

### Quantification of BrdU incorporation into gDNA

HeLa Kyoto cells were cultured for 6 hours after seeding, treated with 40 μM BrdU or DMSO, and cultured for 5 days. Cells were harvested by PBS washing, trypsinization and resuspension in cell culture medium. Pellets were frozen, and genomic DNA extracted using the Monarch gDNA extraction kit (NEB, T3010), and 500 ng of DNA was digested into nucleosides with 5 units of DNA Degradase Plus (Zymo Research, E2020) for 2 hours at 37°C. Deoxyribonucleosides were quantified by RP-LC-MS/MS (Reverse Phase Liquid Chromatography and Tandem Mass Spectrometry) using an Ultimate 3000 HPLC system (Dionex, Thermo Fisher Scientific) directly coupled to a TSQ Altis mass spectrometer (Thermo Fisher Scientific). 1 µl aliquot of acidified sample (1% formic acid) was injected onto a Kinetex C18 column (Phenomenex, 100 Å, 150 x 2.1 mm), employing a 4-minute-long linear gradient from 98% A (1 % acetonitrile, 0.1 % formic acid in water) to 60% B (0.1 % formic acid in acetonitrile) at a flow rate of 80 µl/min and a column temperature of 30°C. Nucleosides were quantified by employing selected reaction monitoring (SRM) with the following transitions: m/z 228 ® m/z 112 (dC); m/z 307 ® m/z 191 and m/z 309 ® m/z 193 (BrdU); m/z 243 ® m/z 127 (dT); m/z 252 ® m/z 136 (dA); m/z 268 ® m/z 152 (dG). A calibration curve of defined standards was used to quantify the relative ratio of BrdU as the molar concentration of BrdU divided by the sum of the molar concentrations of dT and BrdU. Data interpretation was performed using TraceFinder software (Thermo Fisher Scientific).

### Cell synchronization for sister-Pore-C

Cells were synchronized as described (see “Cell Synchronization to G2-phase and DSB Induction”), except 40 μM BrdU (Life Technologies, B23151) was added immediately after aphidicolin release to allow BrdU incorporation during DNA replication. In some samples (Fig. S4), doxycycline treatment and sgRNA transfection were omitted, and cells were arrested for an extended period in G2-phase using RO-3306 (Calbiochem, 217699), following the protocol of (*28*).When applied, RO-3306 was included in all buffers until formaldehyde crosslinking. For Sororin-AID experiments, 500 μM auxin (Sigma-Aldrich, I5148) was added 1 hour prior to aphidicolin washout to induce targeted protein degradation.

### Sister-pore-C

Up to 5 million cells were collected by PBS washing, trypsinization, and resuspension in cell culture medium. Cells were pelleted (1,100 g, 1 minute, RT), washed in 1 ml PBS, then fixed in 1 ml 1% formaldehyde (Thermo Fisher Scientific, 28906) for 10 minutes at RT with rotation. Fixation was quenched by 125 mM glycine for 5 minutes at RT with rotation. Cells were chilled on ice for 10 minutes, pelleted (1,100 g, 1 minute, 4 °C), the supernatant was removed, and the pellet was stored at −70 °C. Cells were resuspended in 500 μl HiC lysis buffer (10 mM Tris-HCl pH 8.0, 10 mM NaCl, 0.2% NP-40, 1× complete EDTA-free protease inhibitor) and incubated for 30 minutes at 4 °C with rotation. After pelleting (2,500 g, 5 minutes, 4 °C), they were washed with 200 μl ice-cold 1.5× rCutSmart Buffer (NEB, B6004), pelleted again, and resuspended in 300 μl ice-cold 1.5× rCutSmart. To mildly denature chromatin, 0.1% SDS was added and samples were incubated for 10 minutes at 65 °C with shaking (300 rpm), followed by cooling on ice. SDS was quenched with 1% Triton-X for 10 minutes on ice. Chromatin digestion was performed with 45 units of NlaIII (NEB, R0125L) in 1× rCutSmart Buffer for 18 hours at 37 °C with intermittent shaking (800 rpm, 30 seconds every 5 minutes). NlaIII was inactivated by heating at 65 °C for 20 minutes (300 rpm). Cells were pelleted (2,500 g, 5 minutes, RT) and resuspended in 250 μl ligation mix (1× T4 ligase buffer, 50 U T4 ligase, 0.1% Triton-X, 100 μg/ml BSA) for 4 hours at RT with rotation. Cells were pelleted (2,500 g, 5 minutes, RT), and genomic DNA was purified using the Monarch gDNA extraction kit (NEB, T3010), including an 18-hour incubation at 65 °C with lysis buffer, Proteinase K, and RNase A to reverse crosslinks. Libraries were prepared using the Oxford Nanopore Technologies (ONT) Ligation Sequencing Kit (LSK109 or LSK110) and sequenced on PromethION R9 flowcells for 72 hours.

### Sister-pore-C data pre-processing

Sister-pore-C samples were pre-processed using a custom Snakemake (*89*) pipeline (https://github.com/gerlichlab/sister-pore-c-snakemake), adapted from the ONT pipeline (https://github.com/nanoporetech/Pore-C-Snakemake) to accommodate sister-chromatid-specific contacts. Key additional steps include generating BrdU calls with DNAscent v2 (*65, 69*), creating label libraries for read-IDs, assigning labeled pairs, splitting pairs into *cis*/*trans*, generating .cool files with *cooler* (*90*) and ICE balancing he combined sister-specific contacts with *cooltools* (*92*).

### Estimating fragment numbers in sister-pore-C datasets and down-sampling

*DuckDB* (*93*) was used to count annotated fragments meeting quality thresholds. The *cooltools* “sample” function (*92*) was then used to down-sample cooler files to match fragment counts.

### Extraction of sample regions

Regions from ICE-corrected Hi-C matrices were extracted using the Python *cooler* API (*90*).

### Sister-pore-C aggregate maps at selected DSB sites

Aggregate maps of Hi-C submatrices around TAD centers (~4 Mb windows) were calculated using *cooltools* (https://github.com/mirnylab/cooltools). Each submatrix was averaged pixel-wise, excluding the main and adjacent diagonals to avoid Hi-C artifacts. Initial exploration of datasets was performed using HiCognition (*81*), while final figures were generated through Jupyter notebooks.

### Sister-pore-C ratio maps at selected DSB sites

Differential contact enrichment was computed as log^2^(DSB-induced / untreated) from Hi-C pileups for cis-sister and trans-sister maps. Contact frequencies were aggregated at target sites with *cooltools* (*92*), averaged for condition-specific pileup matrices, and visualized in *matplotlib* (*82*) and *seaborn* (*83*).

### Sister-pore-C contact probability curves

To investigate whether DSB induction alters chromatin topology and sister chromatid cohesion genome-wide, we compared contact probability [P(s)] curves between untreated and DSB-induced samples. P(s) curves for *cis*- and *trans*-sister contacts were calculated separately using *pairtools* (*94*). Briefly, contacts were binned into 72 geometrically spaced intervals ranging from 1 bp to 1 Gb. The number of contacts in each bin was normalized by the square of the covered base pairs. To correct for differences in sequencing depth, P(s) curves for cis- and trans-sister contacts from the same sample were scaled by a common factor such that the combined area under both curves equaled 1 within genomic separations of 10 kb to 200 Mb.

### Profiling contact density by cross-score

To compare changes in loop abundance (*cis*-sister contacts) and *trans*-sister interactions around target sites upon DSB induction (Fig. 4A-D), we performed cross-score analysis (*95*). Cross-scores from sister-pore-C *cis*- and *trans*-sister contact maps were computed at 10 kb resolution for untreated and DSB conditions, using distance ranges of 30 kb–1 Mb in the *cooltools*.*sandbox*.*cross_score* module. We extracted cross-scores ±2 Mb around DSB sites, with a bin overlapping each DSB site, and computed log^2^(DSB-induced / untreated) to detect changes in *cis*-sister loop abundance and *trans*-sister interactions.

## Data availability

All sister-pore-C and ChIP-seq datasets generated in this study will be deposited in the Gene Expression Omnibus (GEO) upon publication of this manuscript. These datasets are also available upon request. All datasets are based on a custom HeLa Kyoto reference genome (hg19 with SNP correction (*28*)).

## Code availability

All iPython notebooks used for sequencing data analysis will be made available upon publication at https://github.com/gerlichlab/teloni_et_all, including detailed descriptions for each script and the figures generated. The computing environment will be provided as a Docker container: https://hub.docker.com/repository/docker/gerlichlab/gerlich_jupyter (tag ‘version-1.8’), which includes a complete list of software package versions.

## Statistical analysis and sample number

All Hi-C and ChIP-seq datasets presented in this paper are merges of at least two independent replicates. All statistical analyses were performed in Python using *scipy* (*96*), as indicated in the figure legends.

## Published datasets used

Dataset of scsHi-C wild type HeLa cells synchronized to G2 from (*28*) (GEO number: GSM4613674) was used in Fig. 2A, Fig. S2A and Fig. S4A, B

**Fig. S1.**
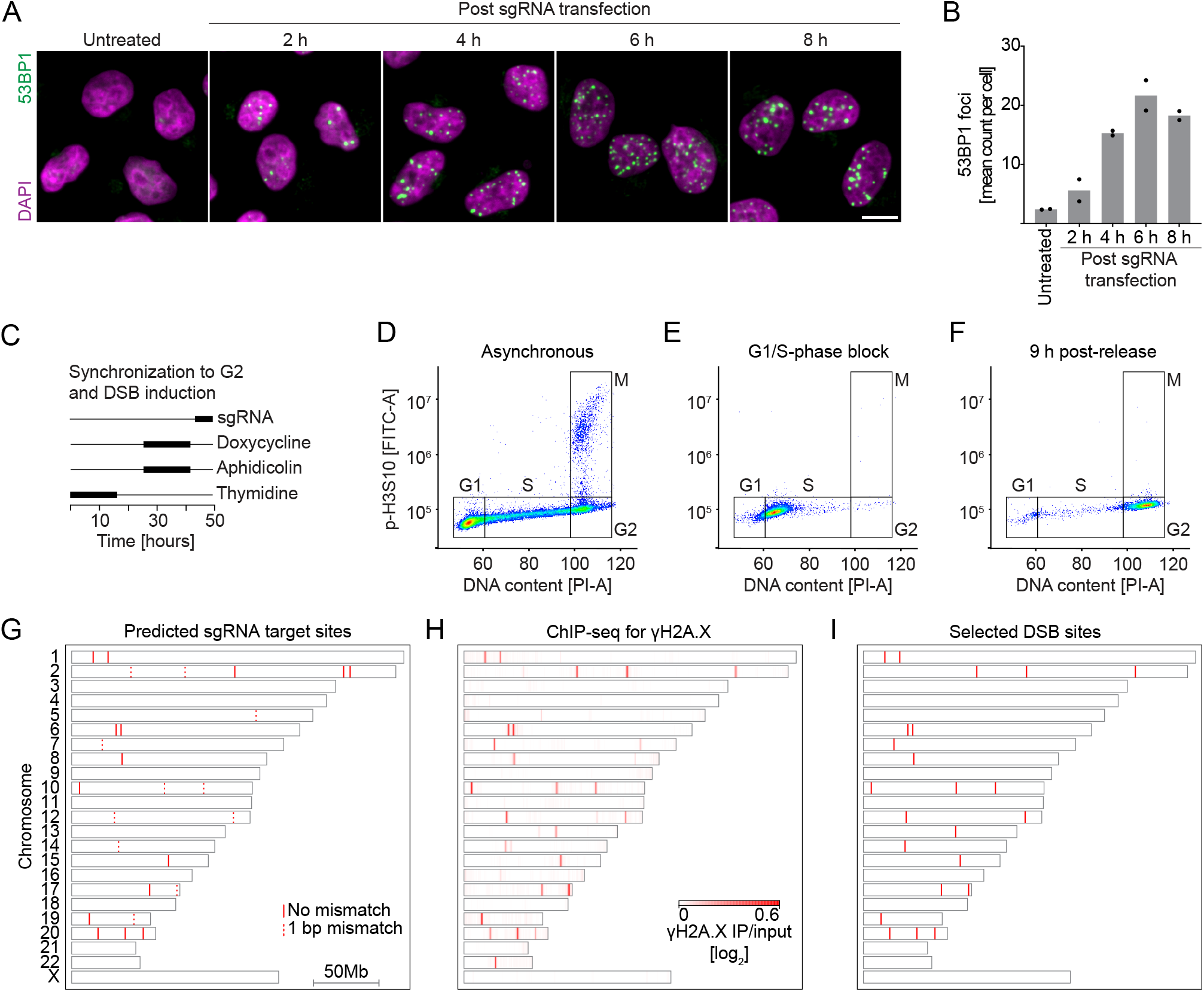
Characterizing the Cas9/sgRNA system for synchronized DSB induction. (**A**) Time course of 53BP1 foci formation following sgRNA transfection in Cas9-expressing HeLa cells. Scale bar, 10 µm. (**B**) Quantification of 53BP1 foci per cell over time (n = 2). (**C**) Schematic of cell synchronization and DSB induction protocol. (**D–F**) Cell cycle distributions (phospho-H3S10 vs. DNA content) in asynchronous (D) or synchronized cells at G1/S (E) or G2 (F). The percentage of G2 cells is 17% (D), 1% (E) and 93% (F). (**G**) Genome-wide map of predicted sgRNA targets (*42*). Red lines, perfect matches; red dotted lines, one-base mismatches. (**H**) ChIP-seq heatmap of γH2A.X in G2-synchronized cells after DSB induction (n = 2). (**I**) Genomic map of the 23 loci selected for subsequent analyses based on γH2A.X enrichment.

**Fig. S2.**
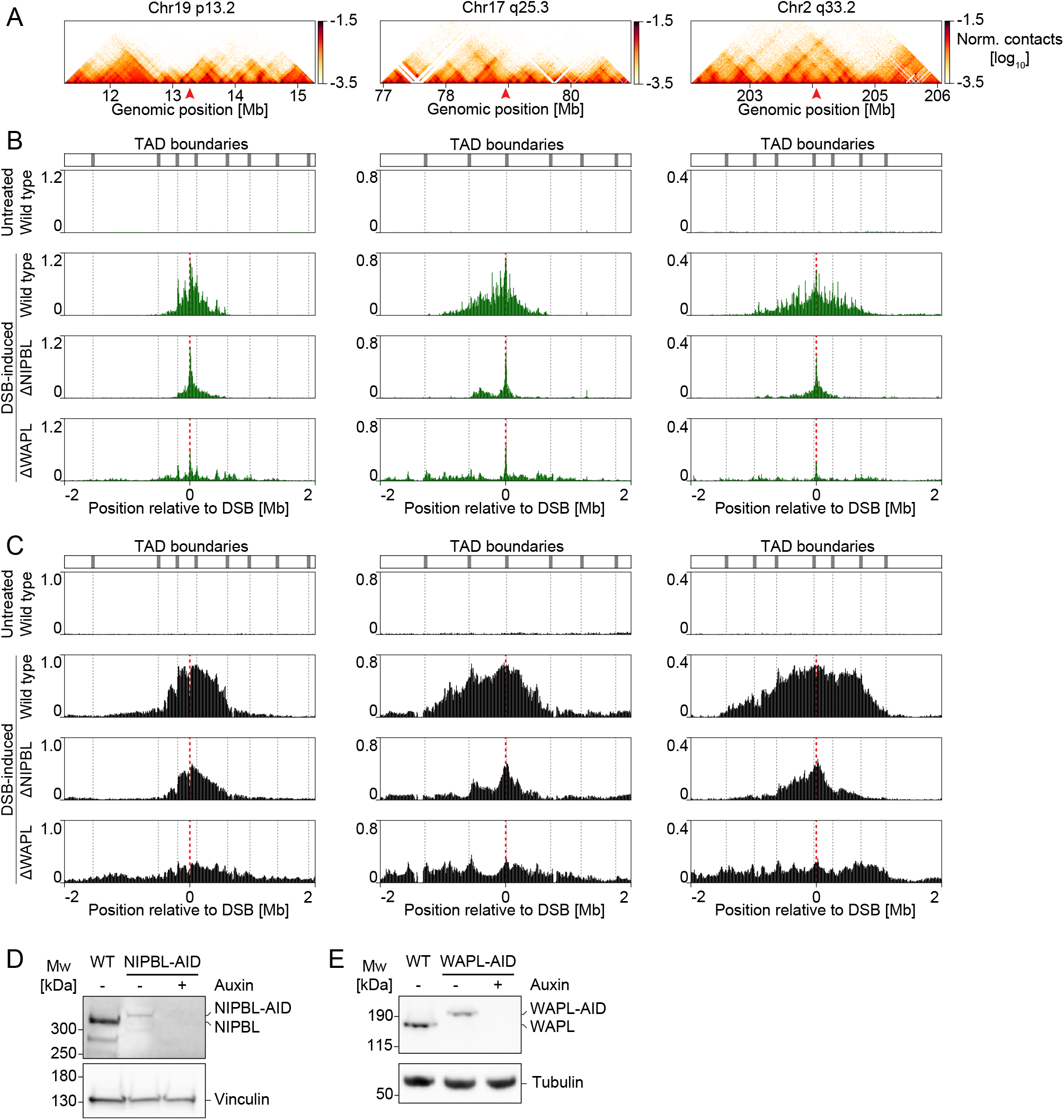
RAD51 and γH2A.X distributions at different sgRNA target sites. (**A**) Hi-C interaction matrices (*28*) at three sgRNA target sites in untreated G2-synchronized cells. Red arrowheads mark the cleavage sites. (**B**) RAD51 ChIP-seq tracks for untreated or DSB-induced G2 cells (wild type or depleted for NIPBL/WAPL). Gray dotted lines indicate TAD boundaries; the red dashed line indicates the cut site. (**C**) γH2A.X ChIP-seq tracks for the same conditions as in (B). All ChIP-seq signals are log^2^(IP/input), merged from n = 2 independent experiments. (**D**) Western blot confirming NIPBL depletion in G2-synchronized cells. (**E**) Western blot confirming WAPL depletion in G2-synchronized cells.

**Fig. S3.**
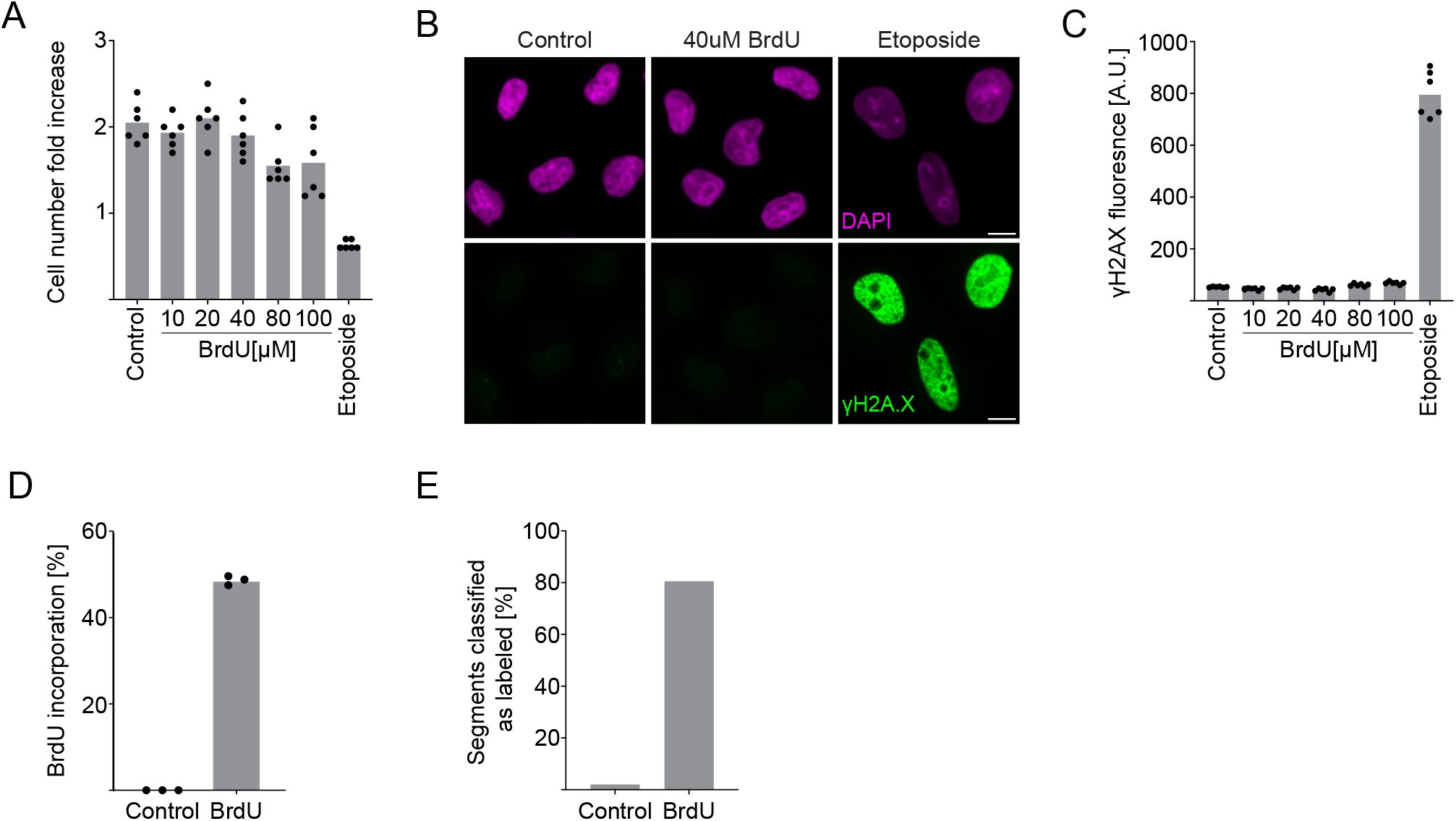
Characterization of BrdU incorporation and its effects on cell viability. (**A**) Fold increase in cell number after 24 h of treatment with the indicated compounds (n = 2). (**B**) Detection of DNA damage (γH2A.X immunofluorescence) after 24 h of the same treatments; scale bar, 10 µm. (**C**) Quantification of γH2A.X fluorescence from (B) (n = 2). (**D**) Mass spectrometry of BrdU incorporation in HeLa cells grown with or without 40 µM BrdU for 5 days, showing BrdU as a fraction of total thymidine (n = 3 technical replicates). (**E**) Detection of BrdU-labeled read segments using DNAscent in sister-pore-C samples with or without 40 µM BrdU as in (D).

**Fig. S4.**
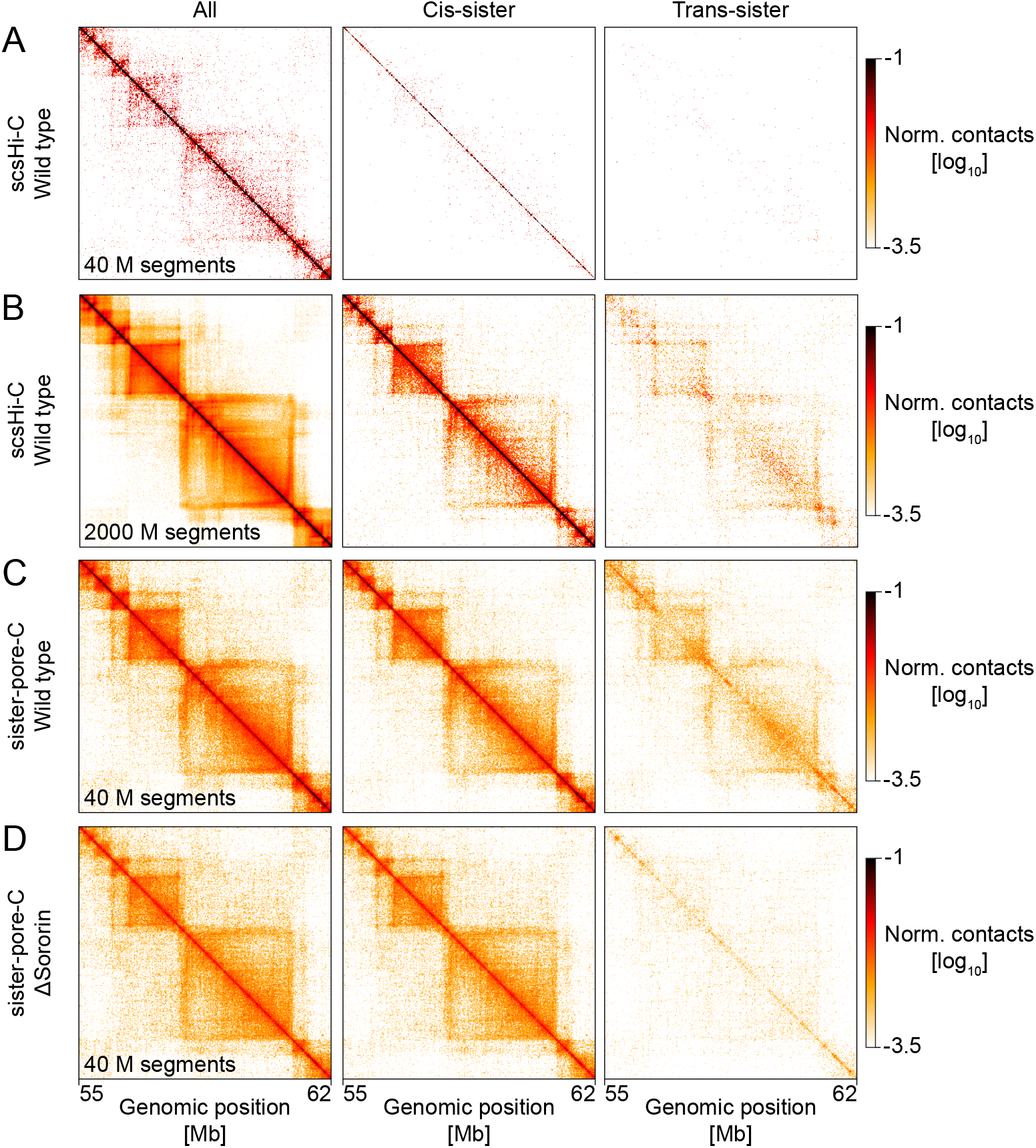
Comparison of sister-pore-C with scsHi-C. (**A, B**) scsHi-C maps of a locus on chr2 in G2-synchronized cells, showing total contacts (left), cis-sister (middle), and trans-sister (right). Data are down-sampled to 40 million segments in (A) and shown at 2 billion reads in (B) (*28*). (**C**) Sister-pore-C maps at the same locus from three merged independent experiments, down-sampled to 40 million segments. (**D**) Sister-pore-C maps in Sororin-depleted cells from two merged independent experiments, also down-sampled to 40 million segments. All data use 20 kb bins. Down-sampling was applied to match the number of sister-specific contacts.

**Fig. S5.**
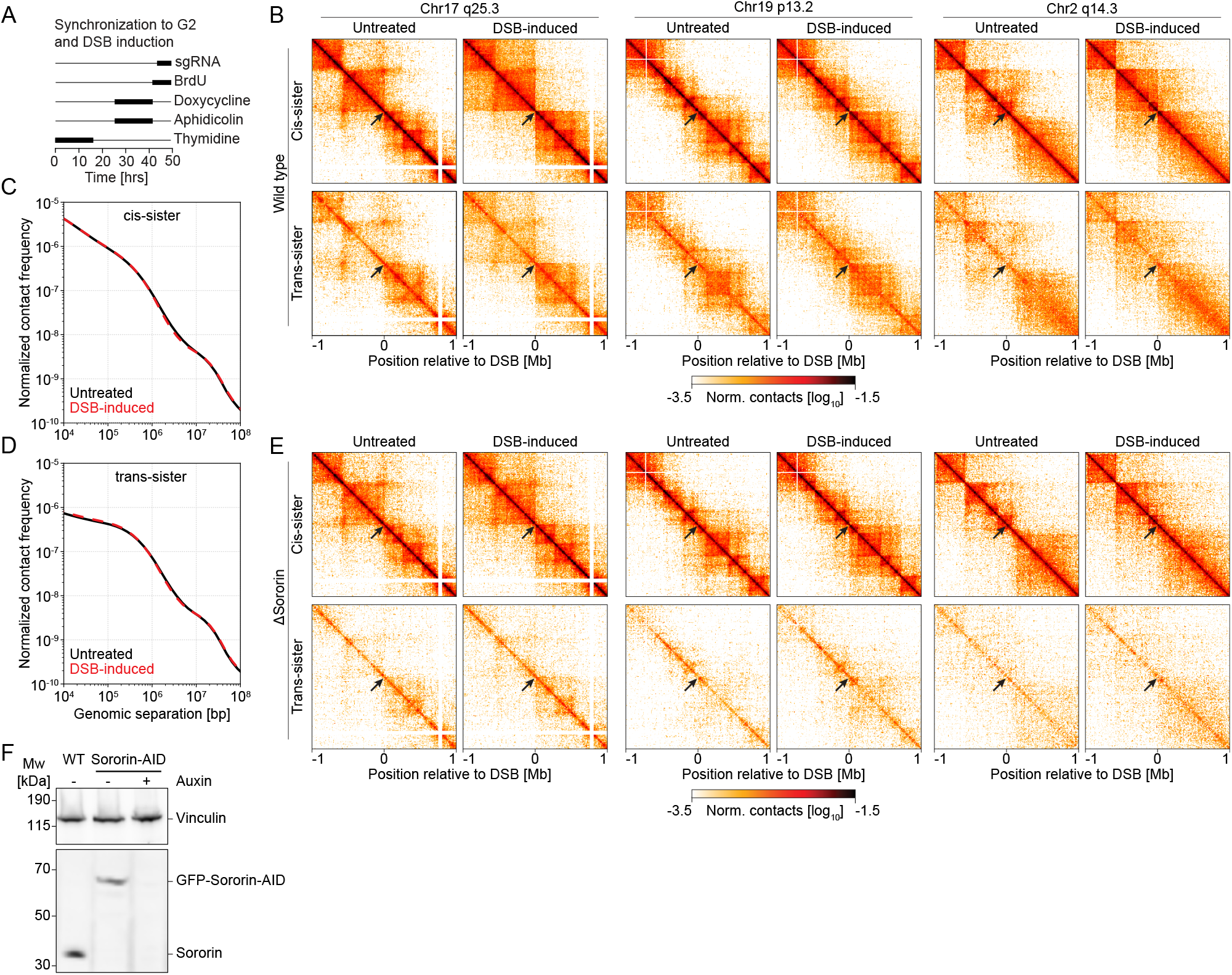
Sister-pore-C analysis at multiple sgRNA target loci. (**A**) Schematic of an updated protocol for G2 synchronization, sgRNA-mediated DSB induction, and sister-pore-C. (**B**) Sister-pore-C matrices (cis-top, trans-bottom) at three loci in untreated or DSB-induced G2 cells. Black arrows mark the DSB sites. Bin size: 10 kb. (**C, D**) Genome-wide contact probability, P(s), over varying distances for cis-(C) or trans-sister (D) contacts in untreated or DSB-induced cells. (**E**) Sister-pore-C matrices at the same loci in Sororin-depleted G2 cells. Data are merged from n = 3 (B–D) and n = 2 (E) independent experiments. (**F**) Western blot confirming Sororin depletion in G2-synchronized cells.

**Supplementary Table 1.** Cell lines used in this study.

| Name | Genotype | Plasmid used | Base cell line used | Resistance marker | Published in |
| --- | --- | --- | --- | --- | --- |
| HeLa Kyoto | WT | n/a | n/a | n/a |  |
| HeLa Kyoto Sororin-AID | EGFP-AID-Sororin<br>TIR1-3xMyc-T2A-Puro (lentiviral integration) | EGFP-AID-Sororin repair template<br>TIR1-3xMyc-T2A PuroLentivirus<br>sgRNA expressing / Cas9D10A expressing to tag Sororin | HeLa Kyoto | Puromycin | 28 |
| HeLa Kyoto Cas9-dox | FKBP12<sup>F36V</sup>-5xGly-3xFLAG-Cas9 (PiggyBac integration) | FKBP12<sup>F36V</sup>-5xGly-3xFLAG-Cas9 (piggyBac with tetracycline response promoter)<br>Super PiggyBac Transposase | HeLa Kyoto | G418 | this study |
| HeLa Kyoto Sororin-AID Cas9-dox | EGFP-AID-Sororin<br>TIR1-3xMyc-T2A-Puro (lentiviral integration)<br>FKBP12<sup>F36V</sup>-5xGly-3xFLAG-Cas9 (PiggyBac integration) | EGFP-AID-Sororin repair template<br>TIR1-3xMyc-T2A PuroLentivirus<br>sgRNA expressing / Cas9D10A expressing to tag Sororin<br>FKBP12<sup>F36V</sup>-5xGly-3xFLAG-Cas9 (piggyBac with tetracycline response promoter)<br>Super PiggyBac Transposase | HeLa Kyoto Sororin-AID (from (28)) | Puromycin, G418 | this study |
| HeLa Kyoto NIPBL-AID Cas9-dox | mEGFP-mAID-NIPBL<br>OstTIR1(F74G)—SNAP-IRES-Blast (integrated into AAVS1 safe harbour locus)<br>FKBP12<sup>F36V</sup>-5xGly-3xFLAG-Cas9 (PiggyBac integration) | mEGFP-mAID-NIPBL repair template<br>OstTIR1F74G-SNAP-IRES-Blast repair template<br>sgRNA / Cas9-human geminin fusion (Cas9-hGem) expressing to tag NIPBL<br>sgRNA / Cas9-human geminin fusion (Cas9-hGem) expressing to integrate OstTIR1F74G<br>FKBP12<sup>F36V</sup>-5xGly-3xFLAG-Cas9 (piggyBac with tetracycline response promoter)<br>Super PiggyBac Transposase | HeLa Kyoto NIPBL-AID (from (27)) | Blasticidin S, G418 | this study |
| HeLa Kyoto WAPL-AID Cas9-dox | Halo-mAID-WAPL<br>SCC1-mEGFP<br>TIR1-3xMyc-T2A-Puro (lentiviral integration)<br>FKBP12<sup>F36V</sup>-5xGly-3xFLAG-Cas9 (PiggyBac integration) | Halo-mAID-WAPL repair template<br>SCC1-mEGFP repair template<br>TIR1-3xMyc-T2A PuroLentivirus<br>sgRNA expressing / Cas9D10A expressing to tag WAPL<br>FKBP12<sup>F36V</sup>-5xGly-3xFLAG-Cas9 (piggyBac with tetracycline response promoter)<br>Super PiggyBac Transposase | HeLa Kyoto WAPL-AID (from (27)) | Puromycin, G418 | this study |

**Supplementary Table 2.** List of sgRNA target sites selected for genomic analysis.

| Chromosome | Genomic location |
| --- | --- |
| chr1 | 16119503 |
| chr1 | 27306872 |
| chr10 | 5904349 |
| chr10 | 99069499 |
| chr10 | 69634394 |
| chr12 | 121354761 |
| chr12 | 32101483 |
| chr13 | 69228918 |
| chr14 | 31260411 |
| chr15 | 72671839 |
| chr17 | 58511366 |
| chr17 | 78950467 |
| chr19 | 13269385 |
| chr2 | 122416332 |
| chr2 | 204055719 |
| chr2 | 85101949 |
| chr20 | 19804371 |
| chr20 | 53691398 |
| chr20 | 40176277 |
| chr6 | 33368117 |
| chr6 | 37059304 |
| chr7 | 22921728 |
| chr8 | 37840790 |

## Notes

### Competing Interest Statement

The authors have declared no competing interest.

